# Electrophysiological evidence for cerebellar involvement in higher-order cognitive processing

**DOI:** 10.1101/294454

**Authors:** Naveen Sendhilnathan, Mulugeta Semework, Michael E. Goldberg, Anna E. Ipata

**Author notes:** These authors contributed equally to this work (Michael E. Goldberg, Anna E. Ipata). Correspondence: Naveen Sendhilnathan, 40 Haven Avenue, Unit 87, 5th floor, Room 561 New York, NY 10032.

## Abstract

Although the cerebellum has been traditionally considered to be exclusively involved in motor control and learning, recent anatomical and clinical studies suggest that it may also have a role in cognition. However, no electrophysiological evidence exists to support this claim. Here we studied the activity of simple spikes of hand-movement related Purkinje cells in the mid-lateral cerebellum when monkeys learned to associate a well-learned right or left-hand movement with one of two visual symbolic cues. The cells had distinctly different discharge patterns between an overtrained symbol-hand association and a novel symbol-hand association although the kinematics of the movement did not change between the two conditions. The activity change was not related to the pattern of the visual symbols, the hand making the movement, the monkeys’ reaction times or the novelty of the visual symbols. We suggest that mid-lateral cerebellum is involved in higher-order cognitive processing related to learning a new visuomotor association.

**One Sentence Summary:** Hand-movement related Purkinje neurons in midlateral cerebellum, which discharge during an overtrained visuomotor association task, change their activity when the monkey has to associate the same movements with new cues, even though the kinematics of the movements do not change.

## Main Text

Historically, the cerebellum has been considered to be exclusively involved in motor control (*1*). Much evidence shows that the cerebellum facilitates motor learning, motor gain adaptation, predictive grip control, timing, and the coordination of motor performance (*2-4*). However, several recent clinical and anatomical studies have challenged this notion. In a classic review, Leiner, Leiner, and Dow argued that the basal ganglia, the thalamus, and the cerebellum all expanded as the primate frontal cortex evolved (*5*) and hypothesized that “Signals from the older part of the dentate nucleus certainly help the frontal motor cortex to effect the skilled manipulation of muscles, and signals from the newest part of the dentate nucleus may help the frontal association cortex to effect the skilled manipulation of information or ideas.” Newer tract-tracing (*6*) and fMRI connectivity (*7*) studies have provided in greater detail evidence that the range of cerebello-cortical connections extends far beyond purely motor areas, encompassing prefrontal areas important in behavior but less so in the more specific aspects of motor control. The presence of cerebellar-cortical-cerebellar anatomical loops suggest that the cerebellum and the cerebral cortex may work in tandem in generating complex cognitive behavior beyond that of fine-tuning movement (*8*). Despite this increasing clinical and anatomical evidence for cerebellum’s role in higher order cognitive functions, no clear electrophysiological evidence exists supporting the claim.

We studied the activity of mid-lateral cerebellar Purkinje neurons in monkeys, during a visuomotor association task. Previous fMRI studies have implicated that the BOLD activity in Crus I and II of the cerebellum increases when normal subjects perform a task that associates an arbitrary visual stimulus with a specific button-press (*9, 10*) suggesting its role in complicated visuomotor association. We trained two macaque monkeys to perform a two-alternative forced-choice discrimination task, where the monkeys associated one of two visual symbols as a cue for a left-hand movement and the other symbol as a cue for a right-hand movement. A trial started when the monkeys placed their hands on two bars after which one of the two symbols appeared on the screen and the monkeys lifted the hand associated with that symbol with a well-learned, stereotypic hand movement to earn a liquid reward. We trained the monkeys to associate a specific pair of symbols (green square and pink square) with specific choices (left and right-hand release, respectively) until their performance was on the average above 95% correct, which we refer to as the overtrained visuomotor association condition. A typical recording session started with the overtrained condition, and on a random trial, we switched the symbols to a pair of non-verbalizable (by humans) fractal stimuli that the monkeys had never seen before. The monkeys then had to learn the correct symbol-hand associations through trial and error (**Fig 1A**). We refer to this as the novel condition.

**Figure 1:**
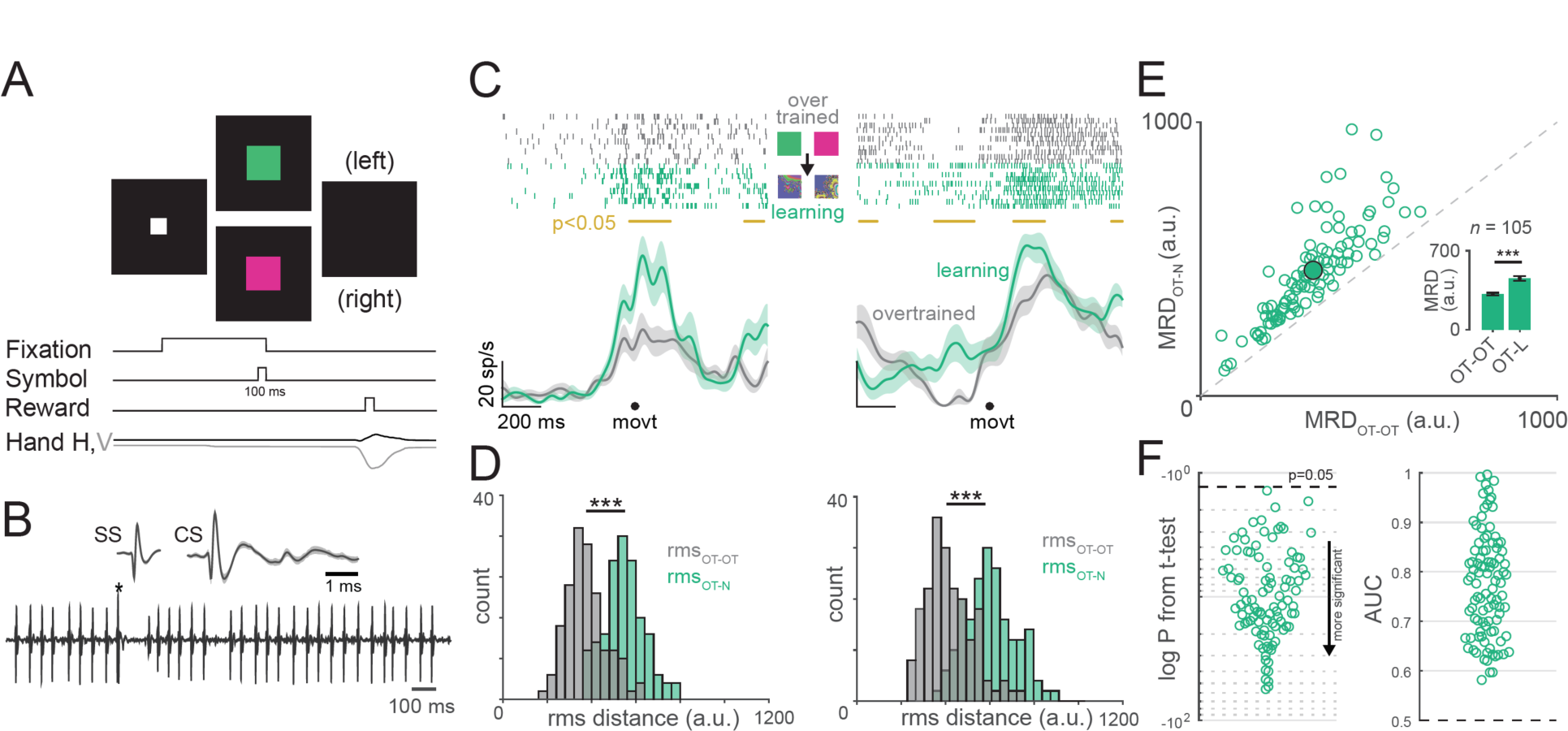
Purkinje neurons showed a change in pattern of neural activity between overtrained and novel visuomotor associations. Two-alternative forced-choice discrimination task: Monkeys were trained to associate a pair of symbols with a pair of choices and report it through hand movement. The correct symbol-choice association is shown for the overtrained condition (Green symbol – left hand; Pink symbol – right hand). Representative recording from a mid-lateral Purkinje neuron showing simple spikes and complex spike (marked by *). Left panel: rasters (top-left) and spike density functions (bottom-left) aligned on hand movement onset, for a representative neuron that increased its activity during the hand movement in overtrained (gray) and novel conditions (green). Gold line in the middle indicates the epochs when the activity between the two conditions were significantly different from each other (p < 0.05 t-test). Right: Same convention as left, but for a representative neuron that decreased its activity after a stimulus onset but also increased its activity during the hand movement in the overtrained condition. Distribution of rms distances between the mean activities of two sets of 10 randomly chosen trials from the last 20 trials before the symbol switch within the overtrained condition (gray) and between the overtrained and novel conditions (green) for the two neurons shown in C. *** means P<0.001; t-test. a.u is arbitrary units. Scatter plot of Mean RMS Distance (MRD) for within overtrained (OT) condition (OT-OT) vs MRD for across overtrained and novel (N) condition (OT-N), obtained from method described in D. Each open circle is a cell and the mean values are shown as filled circles. The inset shows the mean MRD for OT-N and OT-OT conditions. Beeswarm plot of log P values obtained from t-tests (left) and area under the curve of ROC analysis (right) comparing the distribution of the rms distances in the OT-OT condition with that of OT-N condition for all the cells.

We recorded the activity of 105 Purkinje neurons (see Methods and **Fig. S2**) from the left mid-lateral cerebellum of two monkeys (80 from monkey B; **Fig S1A** and 25 from monkey S; **Fig S1B**). Of these, 88 neurons increased their firing during bar-release hand movement (**Fig 1C**) and 32 of these neurons showed a stimulus related decrease as well as a hand related increase in activity. These neurons did not have a preference for specific visual symbols (**Fig S3A, B;** P = 0.3332, one-sample t-test) and responded during either hand movement (**Fig S3C, D;** P = 0.5790, one-sample t-test).

The activity of Purkinje cells changed dramatically at the switch from overtrained to novel visuomotor association (*P* < 10^−8^ Mann-Whitney U-test; *P* < 10^−21^; two sample KS test) in multiple epochs of the trial (**Fig 1C,** indicated by gold lines on the top). To quantify this change in activity, first, we randomly sampled 10 trials each from the last 20 trials in the overtrained condition and the first 20 trials in the novel condition and calculated the root mean squared (rms) distance between the mean activities. We repeated this process 250 times to obtain a distribution of rms distances that compared the extent of change in across-condition activity profile in the novel condition from the activity profile in the overtrained condition (green histograms in **Fig 1D**). To compare this distribution with a control null-distribution, we randomly sampled 10 trials twice without replacement from the overtrained condition and repeated the same analysis to obtain another distribution of rms distances (gray histograms in **Fig 1D**) to obtain an estimate of variability of within-condition. Across the population, the within-condition rms values were significantly lower than the across-condition rms values indicating that the change in activity at the switch to novel condition could not be explained by the variability in activity in the overtrained condition. Every cell we studied significantly changed its activity significantly between the two conditions (**Fig 1F** left panel: log P values from t-test; right panel: area under the curve (AUC) from ROC analysis between the two distributions). See **Fig S5** for validity of this method.

Because midlateral cerebellar Purkinje neurons change their activity with changes in movement (*11*), we then asked if the changes in neural activity could be attributed to any changes in motor kinematics. We only tracked the monkeys’ right-hand movement in space and time through a high-speed camera because the neurons had similar response with either hand movement (**Fig S3D**). The monkeys performed very stereotypic and consistent hand movements, releasing the bars, to indicate their choice. Although the hand movement did not change at the visuomotor association switch (**Fig. 2A,** Representative session: **Fig S7A-D;** Population: **Fig S7E, F**; *P* = 0.3822, t-test), the neural activity changed markedly (**Fig 2B, C;** *P* < 10^−3^, Wilcoxon rank sum test). Therefore, we could not attribute the change in neural activity to a change in the hand movement. Although the motor kinematics were quite different between the two monkeys (**Fig S6**), in neither monkey did the movement change at the visuomotor association switch.

**Figure 2.**
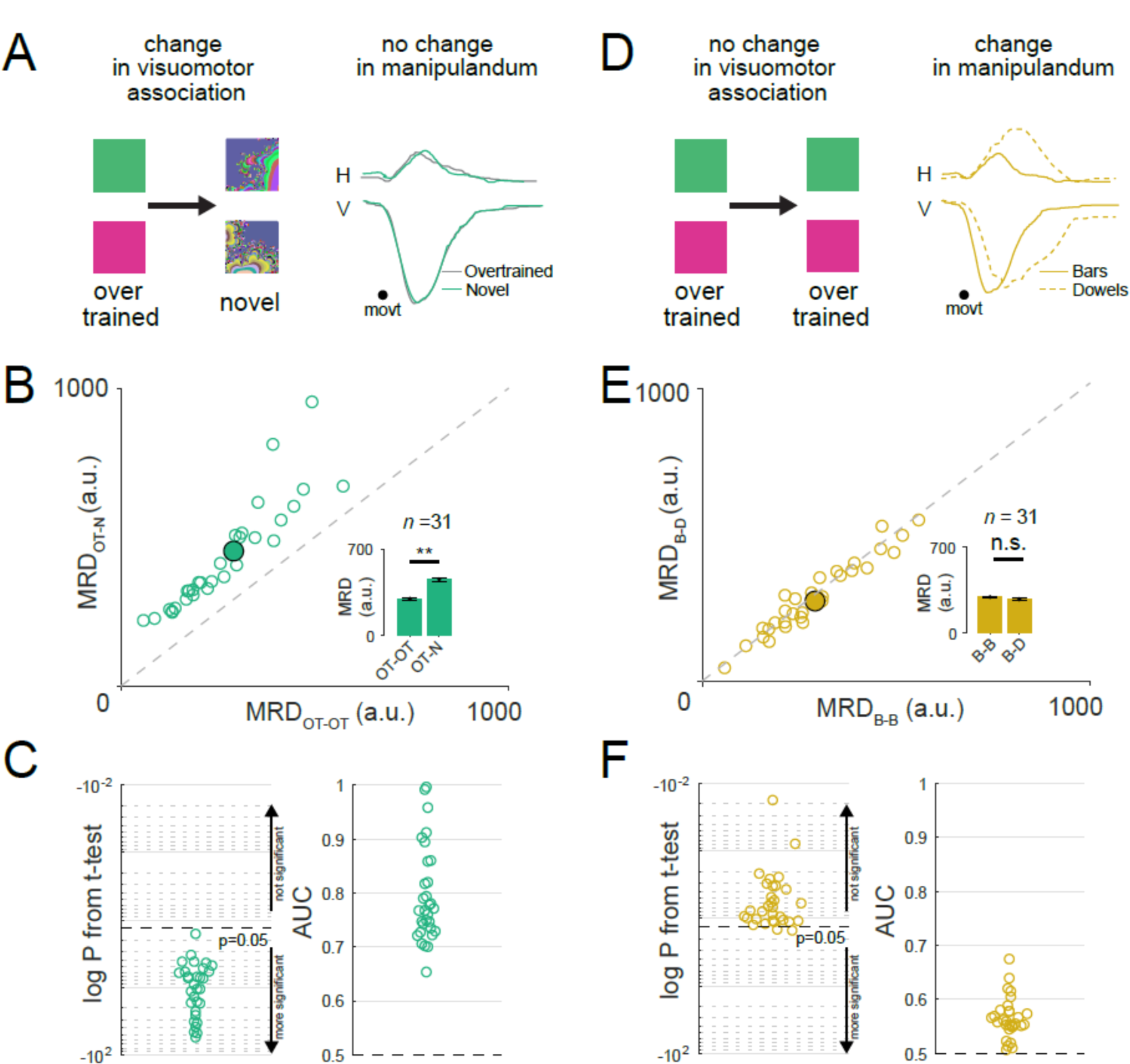
Mid-lateral Purkinje neurons do not encode specific motor kinematics. Left panel: Task condition with change in visuomotor association (left) but no change in manipulanda (right). Right panel: Horizontal and vertical hand movement traces for overtrained (grey) and novel (green) conditions. Scatter plot of MRD for within overtrained condition (OT-OT) vs MRD for across overtrained and novel condition (OT-N) for the same neurons shown in 2B. Note that the neurons shown here are a subset of those shown in Fig 1E. Same convention as Fig 1E. ** means P<0.01; t-test. Beeswarm plot of log P values obtained from t-tests (left) and area under the curve of ROC analysis (right) comparing the distribution of the MRDs in the OT-OT condition with those of OT-N condition. Same convention as Fig 1F. Left panel: Task condition with no change in visuomotor association (left) but a change in manipulanda (right). Right panel: Horizontal and vertical hand movement traces for flat bars (solid line) and dowel (dotted line) conditions. Scatter plot of MRD for within bars condition (B-B) vs MRD for across bars and dowels condition (B-D). Same convention as Fig 1E. n.s. means not significant; t-test. Beeswarm plot of log P values obtained from t-tests (left) and area under the curve of ROC analysis (right) comparing the distribution of the MRDs in the B-B condition with that of B-D condition. Same convention as Fig 1F.

To test if these neurons were truly movement invariant, we altered the movements associated with manipulanda release while keeping the visuomotor association constant. Because the monkeys were highly habituated to the bar manipulanda, we switched the manipulanda to a pair of dowels (cylindrical rods, upon which the monkeys were seldom trained for a long time), on a randomly chosen trial, while keeping the overtrained visuomotor association the same. Although the kinematics of the movement changed markedly (**Fig. 2D,** Representative session: **Fig S7G-J;** Population: **Fig S7K;** *P* < 10^−48^, t-test), the neural activity did not change significantly (**Fig 2E, F;** *P* = 0.5563, t-test). These two experiments suggest that the mid-lateral cerebellum is involved in establishing visuomotor associations, a cognitive process, rather than in adjusting or specifying the kinematics of movement. Nevertheless, we cannot exclude the possibility that other body movements unaccounted for by our tracking and analyses, including minor differences in grip, movement of digits, gross arm movement etc., could potentially contribute to the observed changes in neural activity.

Although the kinematics of the movement did not change, the reaction time increased significantly in most sessions at the visuomotor association switch (**Fig 3A;** *P* < 10^−14^, Wilcoxon rank sum test). Because monkeys with cerebellar lesions have longer reaction times in fast extensions and flexions (*12*), we wanted to investigate if the changes in neural activity at the visuomotor association switch could be merely attributed to changes in reaction time. We found that in 24/105 (23%) of sessions the reaction time did not significantly change after the visuomotor association switch (**Fig 3B;** *P* = 0.8151, Wilcoxon rank sum test) even though the monkeys’ performance decreased significantly (**Fig 3B;** *P* < 10^−7^, Mann-Whitney U-test). Additionally, we used two different stimulus presentation durations (see methods) but the stimulus duration did not affect the reaction time of the monkey. Nonetheless, the neural activity changed significantly even when the reaction time did not change, and there was no difference in the neural activity between sessions where the reaction time changed and sections where it did not (**Fig 3C, D**; RT change case: *P* < 10^−6^, Mann-Whitney U-test; no RT change case: *P* < 10^−3^, t-test). This suggests that the change in reaction time at the visuomotor association switch did not contribute to the change in neural activity.

**Figure 3:**
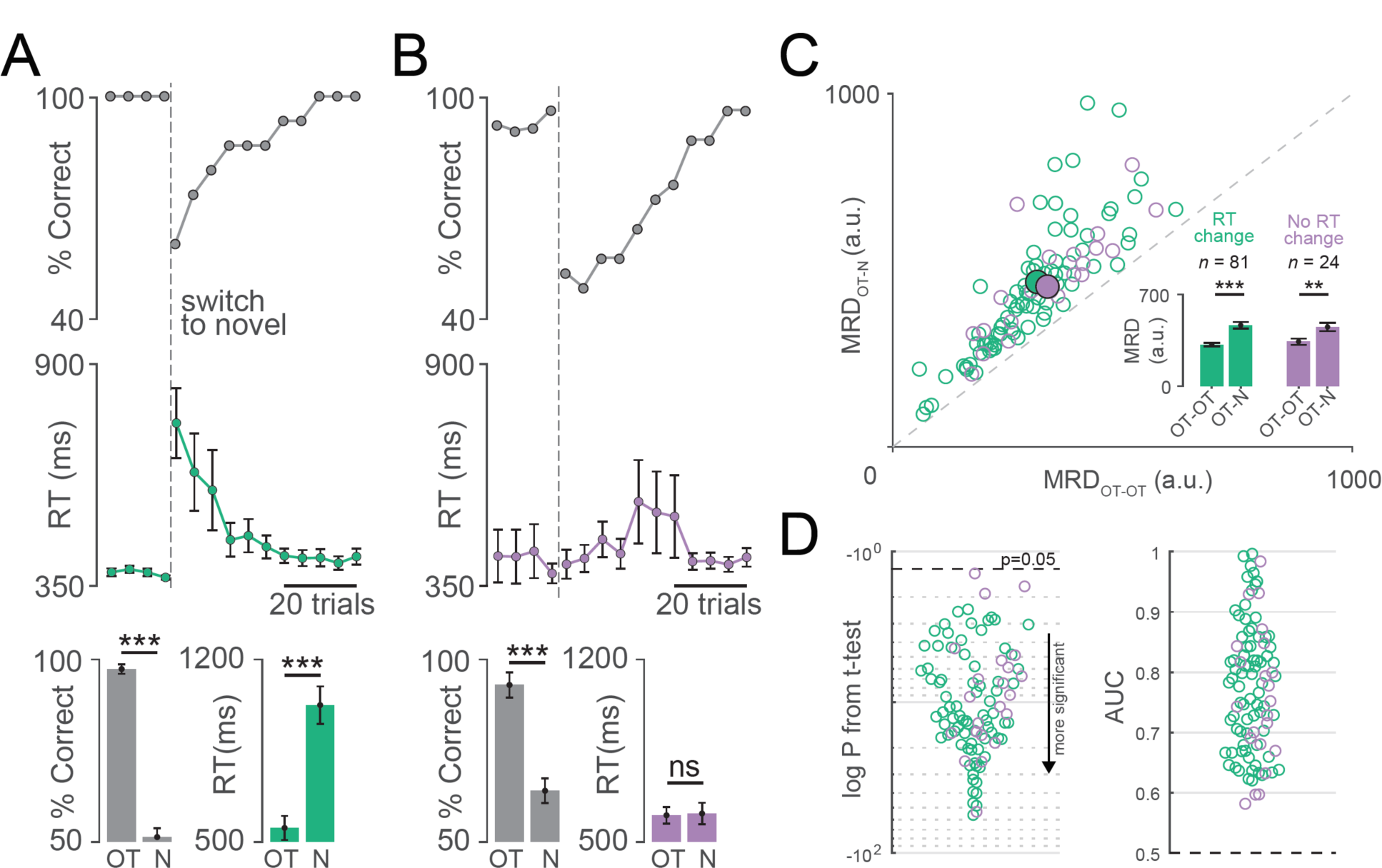
Neural activity changes are independent of reaction time. Top: Percent of correct trials plotted as a function of trial number relative to the switch to novel visuomotor association. Middle: Reaction times for the same trials. Error bars show the standard error of the mean. Bottom: Mean percent correct (left) and reaction time (right) in the overtrained (OT) and novel (N) conditions for all the sessions with changes in reaction time. *** means P<0.001; n.s. means not significant, Mann-Whitney U-test. Percent correct(top), reaction time(middle), and session averages(bottom) for sessions in which the manual reaction time did not change after the switch to novel visuomotor association. Same convention as A. Same data as in Fig 1E but separated into sessions with RT change (green) and no RT change (violet). Same data as in Fig 1F but separated into sessions with RT change (green) and no RT change (violet).

Finally, we investigated if the change in neural activity were due to the novelty of the fractal symbols rather than a change in visuomotor association. On 24 sessions, after the monkeys had learned the novel associations, we reversed the symbol-hand associations and the monkeys had to relearn the associations. Here, there was no change in the symbols but the monkeys had to learn a new visuomotor association nevertheless (**Fig 4A**; *P* < 10^−7^, Mann-Whitney U-test). We again observed that the neurons showed a change in activity (**Fig 4B, C**; *P* < 10^−5^, t-test; *P* < 10^−16^; two sample KS test). This suggests that the change in neural activity could be observed without a change in the symbols if the visuomotor association changed.

**Figure 4.**
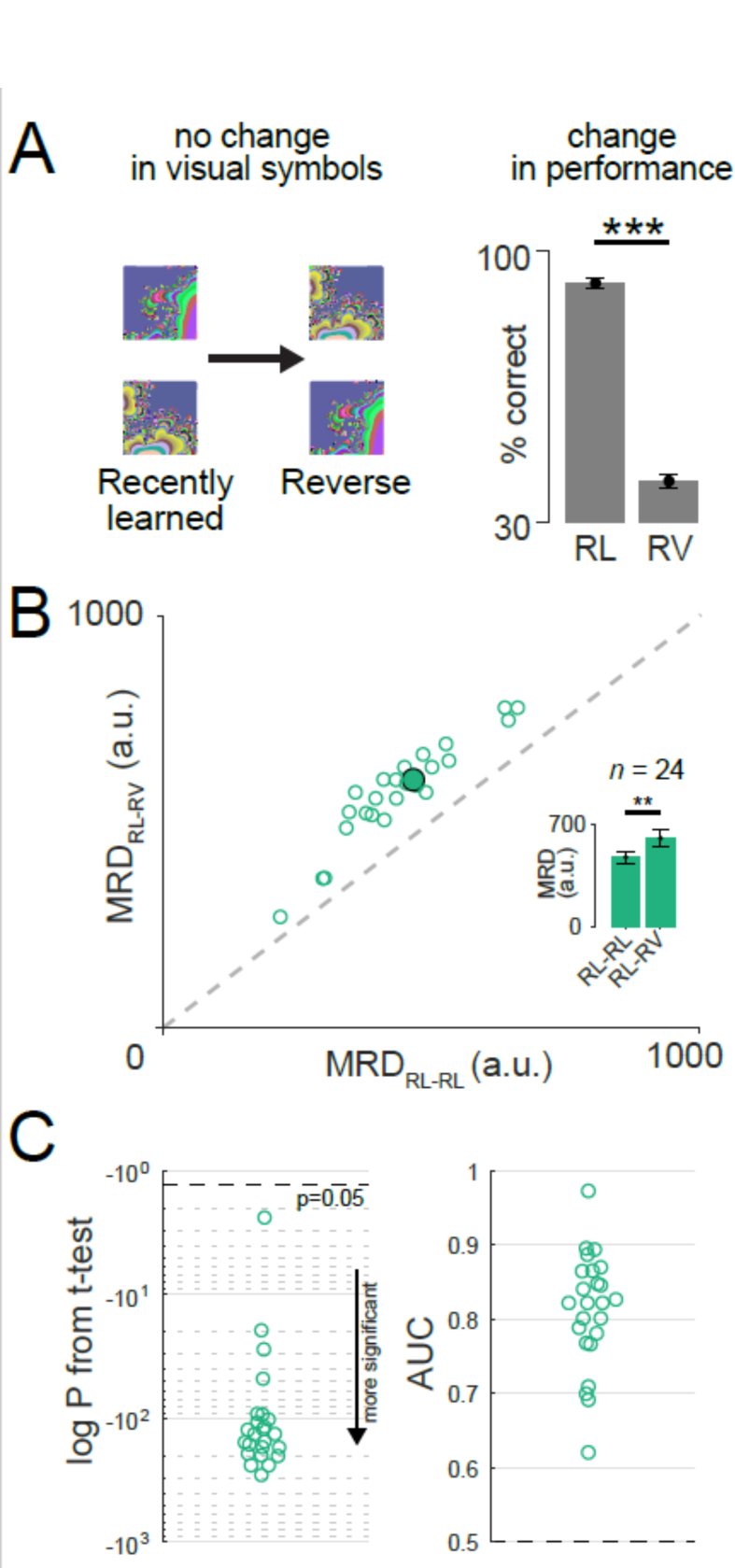
Neural activity changes are independent of novelty of the symbols. Left: Changing the significance of previously seen symbols from recently well-learned to reversal learning condition but with new learning. Right: Mean behavioral performance in the recently learned (RL) and the reversal (RV) conditions. *** means P<0.001, Mann-Whitney U-test. Scatter plot of MRD for within recently learned condition (RL-RL) vs MRD for across recently learned and reverse condition (RL-RV). Same convention as Fig 1E. ** means P<0.01, t-test. Bee swarm plot of log P values obtained from t-tests (left) and Area under the curve of ROC analysis (right) comparing the distribution of the MRDs in the RL-RL condition with that of RL-RV condition. Same convention as Fig 1F.

These results suggest that the mid-lateral cerebellum participates in visuomotor association and not the specification of the motor kinematics dictated by the task. Cerebellar activity changed at the visuomotor association switch where the movement did not change (**Fig 1, S7**), and not when the required movement changed without a concurrent change in the visuomotor association (**Fig 2, S7**). The activity change was independent of the hand used (**Fig S3**), the symbol that evoked the activity (**Fig S3**), the monkey’s reaction time (**Fig 3**) and the novelty of the symbols used (**Fig 4**). We suggest that this region of the cerebellum is important in learning a new visual association for a simple movement.

Strick et. al. (*13*) described two distinct, almost functionally disparate, anatomical regions of the cerebellum: One having connections to the motor regions of the cortex and the other to the non-motor or cognitive regions of the cortex. The area that we recorded from, in this study, is predominantly a hand movement region in what seems like the putative cognitive region of the cerebellum. Our results show that although these neurons might encode certain aspects of the movement when the monkeys performed an overtrained visuomotor association, they change their activity when the monkeys must learn a new visuomotor association even though the movement does not change.

Changing neural activity related to novel visuomotor association has been previously reported in the monkey prefrontal cortex and basal ganglia (*14, 15*), areas shown to be within a loop shared by the cerebellum (*16*) as well as other areas. However, a similar function has not been previously found in cerebellar neurons in the monkey. Our result suggests that mid-lateral cerebellar Purkinje neurons have properties similar to the prefrontal neurons involved in higher order cognitive processing. We show the first electrophysiological evidence in the monkey for the participation of cerebellum in higher order cognitive processing and thus provide a major departure from our current understanding of the role of cerebellar Purkinje neurons.

## Acknowledgments

This work would not have been possible without the wonderful contributions of John Caban for designing and building the manipulandum changer and the video mount; Glen Duncan for creative electronic assistance, Matthew Hasday for superb machining, Dr. Girma Asfaw, Dr. Moshe Shalev for animal care, Dr. Vincent Prevosto for helping us install the CED1401 and interfacing it with REX, and Lisa Kennelly and Holly Cline for facilitating everything. This work was supported by the Keck, Zegar Family, and Dana Foundations and the National Eye Institute (R24 EY-015634, R21 EY-017938, R21 EY-020631, R01 EY-017039, P30 EY-019007, and R01 EY-014978 to M. E. Goldberg, PI.

---

**Figure S1:**
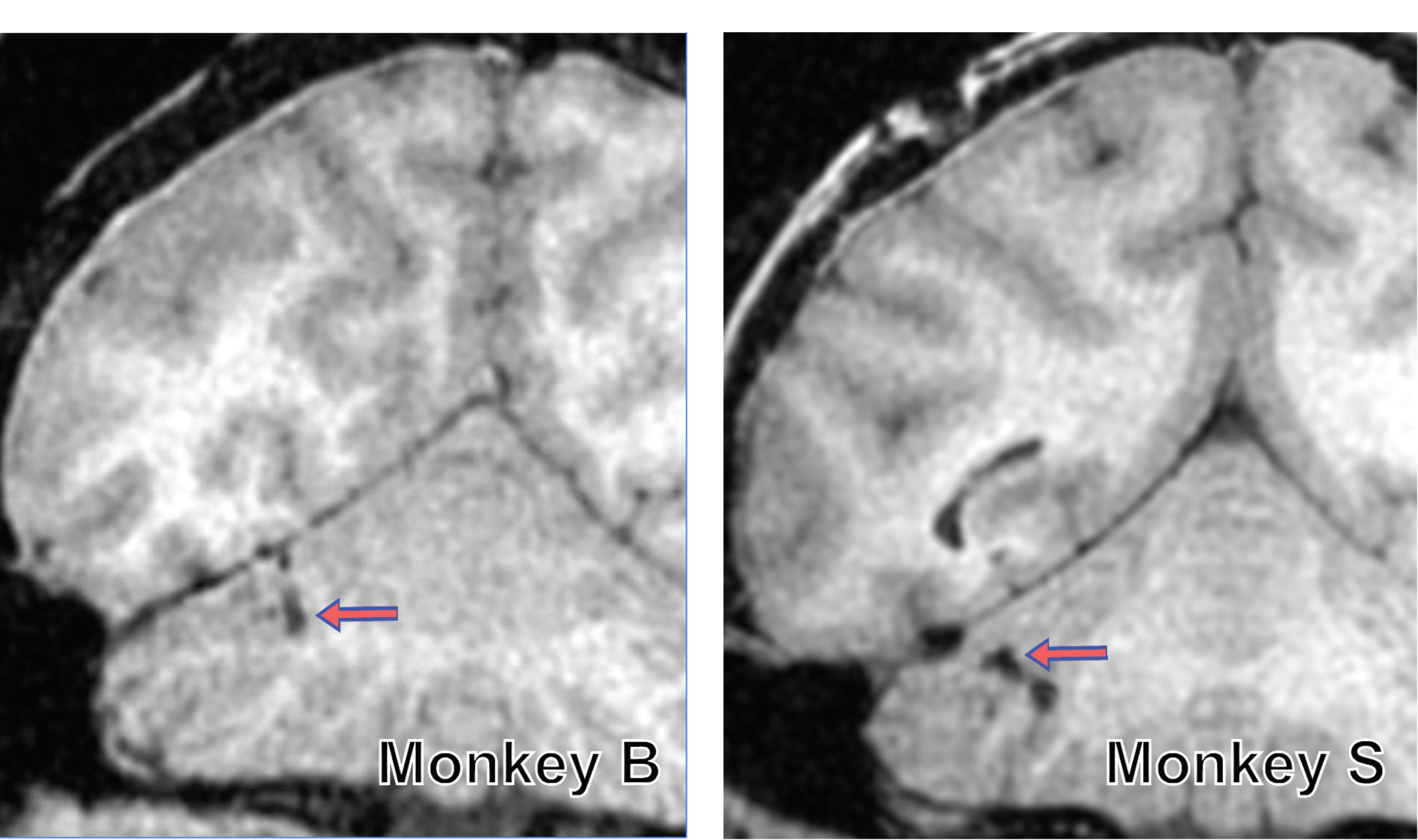
MRI of recording sites from mid-lateral cerebellum. MRI images of the monkeys’ brain showing the cerebellum for monkeys B (left), and S (right) showing the site of recording (red arrow) in the mid-lateral cerebellum.

**Figure S2.**
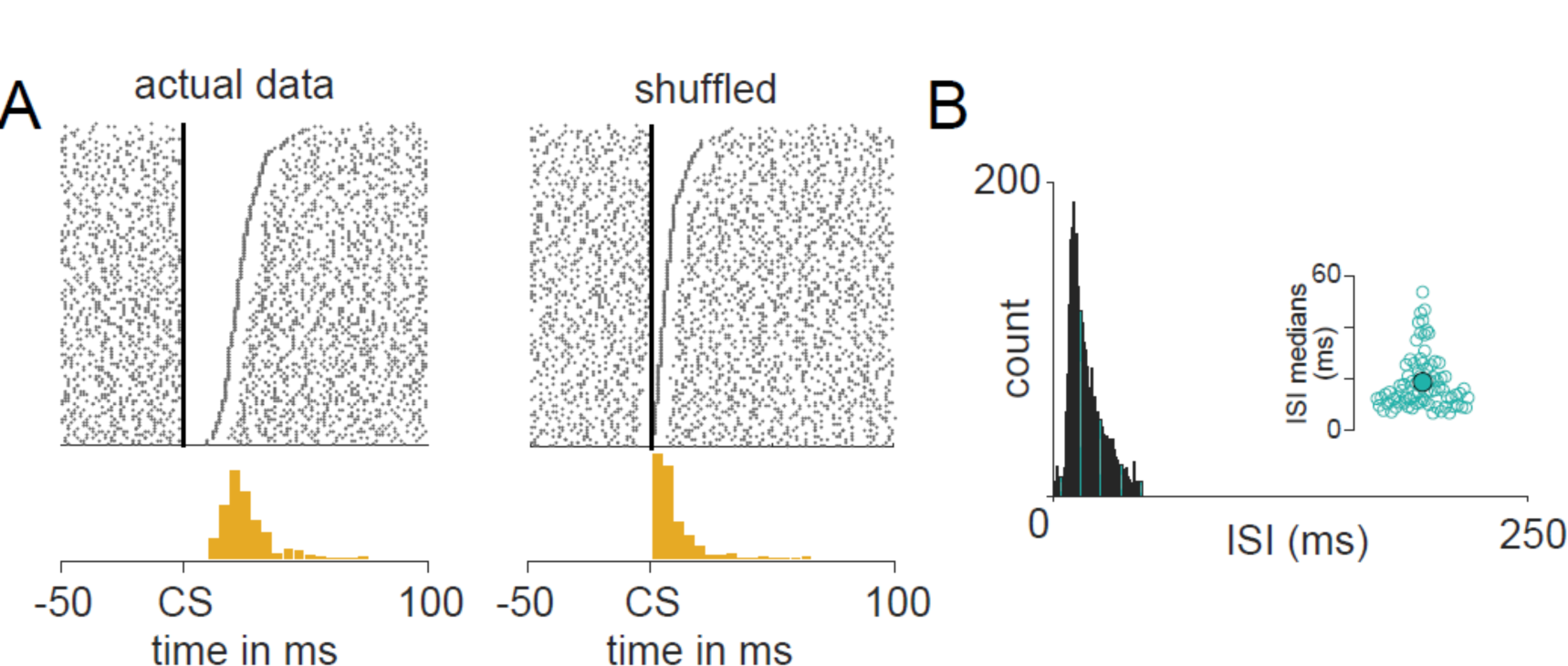
Identification of Purkinje neurons. **A.** Top left: Simple spikes aligned to the preceding complex spikes. Bottom left: Histogram of time of occurrence of the first simple spike after every complex spike. Note that the distribution has a lag from 0 ms. Top right: Same data from top left, but with shuffled simple spike timings. Bottom right: Same as bottom left, but for data in top right. **B.** Interspike Interval distribution for simple spikes from a representative neuron. Inset shows the median of the interspike interval (ISI) for all the recorded Purkinje neurons.

**Figure S3.**
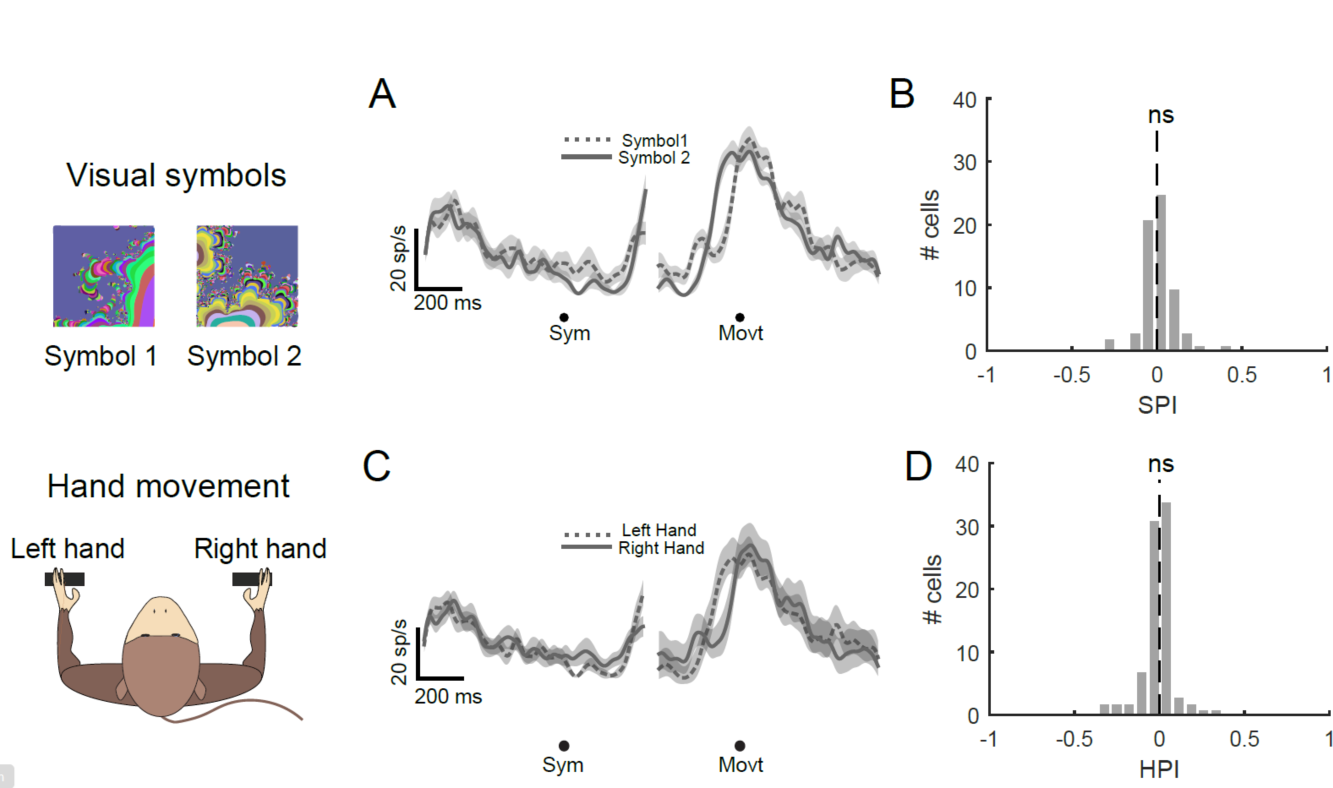
Purkinje neuron simple spikes were neither symbol selective nor hand selective. **A.** A representative neuron’s activity in the overtrained condition for symbol1 (dotted line) and symbol2 (solid line) aligned to symbol onset and movement onset. **B.** Histogram of Symbol Preference Index (SPI) calculated for all neurons in the overtrained condition. n.s. means not significantly different from 0; t-test. This means the neural activity in the overtrained condition has no symbol preference. **C.** Same as A., but for left vs right hand. **D.** Same as B., but for hand preference index (HPI). n.s. means not significantly different from 0; t-test. This means the neural activity in the overtrained condition has no hand preference.

**Figure S4.**
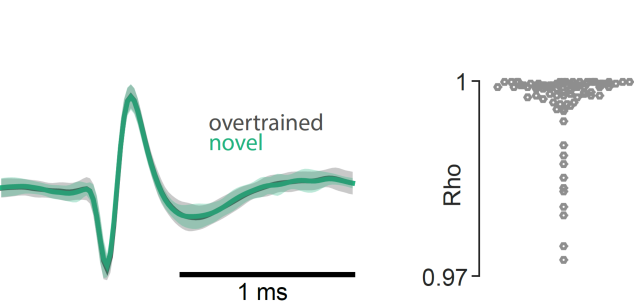
Electrode drift does not explain the change in pattern of neural activity. Left: average spike waveforms for overtrained condition (grey) superimposed on novel condition (blue) for one session. Right: beeswarm plot of correlation between the spike waveforms on overtrained and novel conditions for all sessions. Each circle is a neuron.

**Figure S5.**
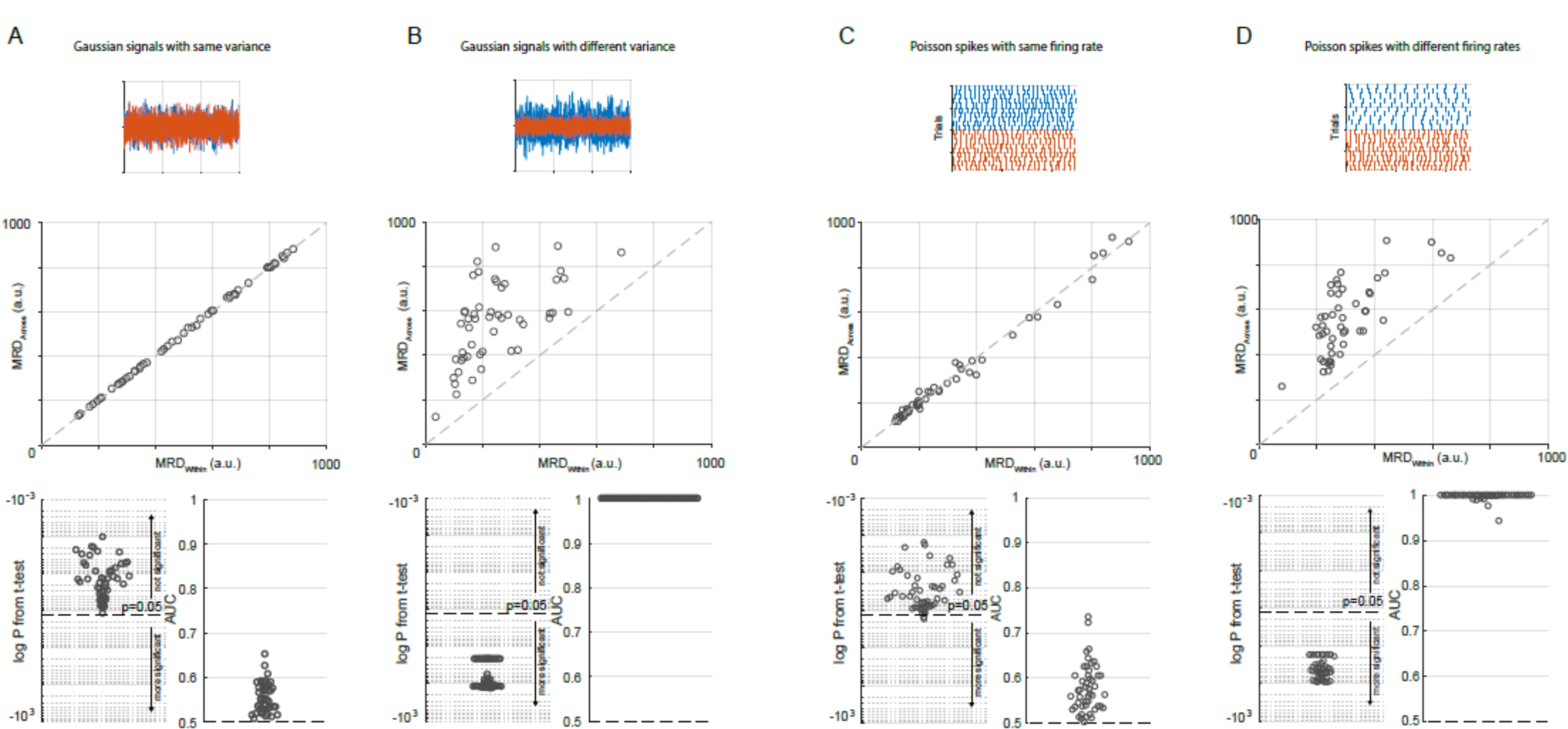
Validity of the MRD method. **A.** MRD Method applied to 50 pairs of Gaussian signals with 0 mean and same variance. Top panel shows one such pair. Middle panel shows the Mean RMS Distance (MRD) calculated across group plotted against that calculated within the first group. Bottom left: log P from t-test between the MRD_across_ and MRD_within_. Bottom right: Area under the curve (AUC) from a ROC analysis comparing the distribution of MRD_across_ and MRD_within_. **B.** Same as A, but for 50 pairs of Gaussian signals with 0 mean and different variances. **C.** Same as A, but for 50 pairs of mean spike density functions obtained from Poisson spike trains with same mean and variance (λ). **D.** Same as C, but for 50 pairs of mean spike density functions obtained from Poisson spike trains with different means and variances.

**Figure S6.**
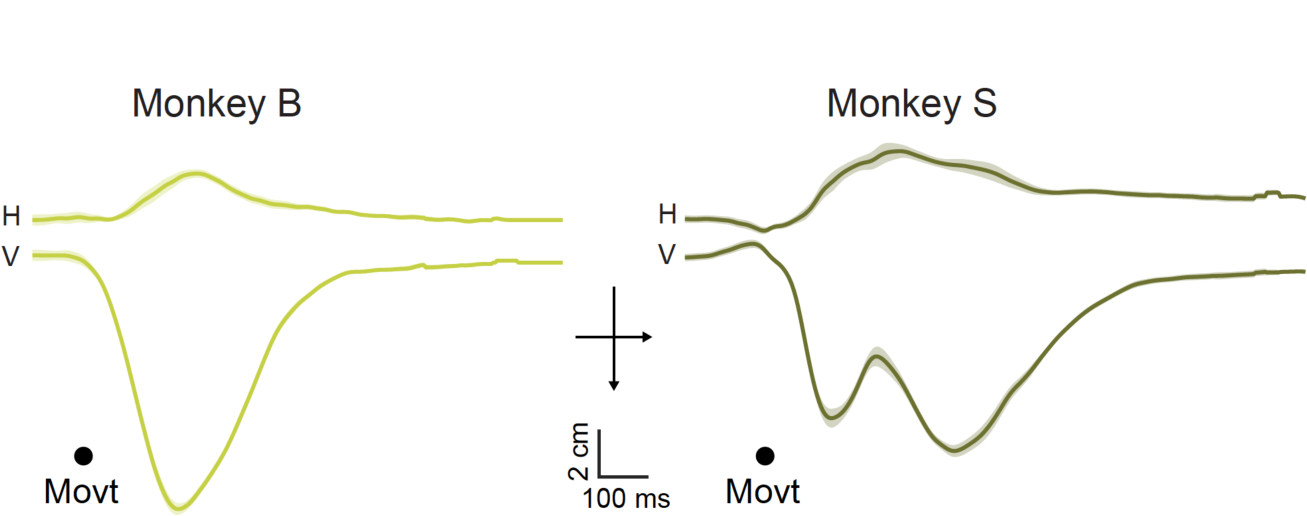
The learned hand movements were precise but distinct for different monkeys. Horizontal (H) and Vertical (V) mean hand trajectories, aligned to the time of bar release (movt) for both monkeys during overtrained condition.

**Figure S7.**
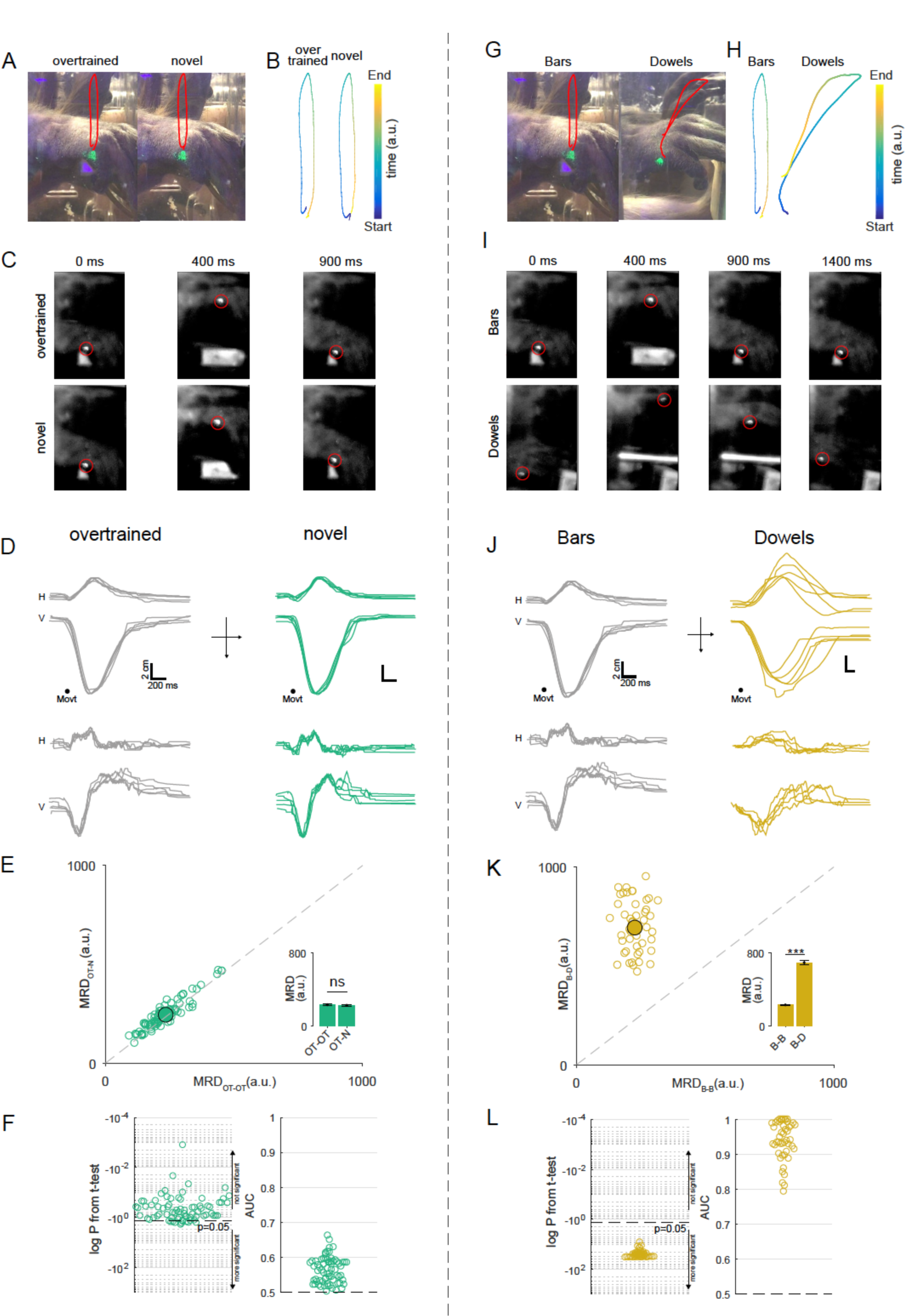
Motor kinematics. **A.** Schematic of the hand movement trajectory during bar release in the overtrained (left) and novel conditions (right). The red trace maps the position of the green fluorescent marker in space through the hand movement. **B.** Position of the fluorescent marker in space and time (color indicates the time as in the colorbar inset). **C.** Snapshots from high-frame rate movies showing the monkey’s hand movement trajectory at three time points (0, 400 and 900 ms from the start of movement) for overtrained (top three panels) and novel (bottom three panels) conditions. Red circle marks the fluorescent marker. **D.** Top panel: Horizontal (H) and Vertical (V) hand trajectories for five continuous trials aligned on movement onset for overtrained condition. Bottom panel shows the H and V velocities for the same five trials. **E.** Scatter plot of Mean RMS Distance (MRD) for within overtrained (OT) condition (OT-OT) vs MRD for across overtrained and novel (N) condition (OT-N), for hand movement trajectories. Same convention as Fig 1E. **F.** Beeswarm plot of log *P* values obtained from t-tests (left) and area under the curve of ROC analysis (right) comparing the distribution of the rms distances in the OT-OT condition with that of OT-N condition for all the hand movements. Same convention as Fig 1F. **G.** Schematic of the hand movement trajectory during bar release (left) and the dowel release (right) conditions. The red trace maps the position of the green fluorescent marker in space through the hand movement. **H.** Position of the fluorescent marker in space and time (color indicates the time as in the color bar inset). **I.** Snapshots from high-frame rate movies showing the monkey’s hand movement trajectory at four time points (0, 400 and 900 and 1400 ms from the start of movement) for bar release (top three panels) and dowel release (bottom three panels) conditions. Red circle marks the fluorescent marker. **J.** Top panel: Horizontal (H) and Vertical (V) hand trajectories for five continuous trials with, bar release, aligned on movement onset. Bottom panel shows the H and V velocities for the same five trials. **K.** Scatter plot of MRD for within bars (B) condition (B-B) vs MRD for across bars and dowels (D) condition (B-D), for hand movement trajectories. Same convention as Fig 1E. **L.** Beeswarm plot of log *P* values obtained from t-tests (left) and area under the curve of ROC analysis (right) comparing the distribution of the rms distances in the B-B condition with that of B-D condition for all the hand movements. Same convention as Fig 1F.

## Methods

### Animal subjects and surgery

We used two male adult rhesus monkeys (*Macaca mulatta*), B and S, weighing 10-11 kg each, for the experiments. All experimental protocols were approved by the Animal Care and Use Committees at Columbia University and the New York State Psychiatric Institute, and complied with the guidelines established by the Public Health Service Guide for the Care and Use of Laboratory Animals. We located the cerebellum in each monkey using a Bravo volume scan using a GE 3T magnet, with a tungsten recording electrode placed at a position from which we had recorded hand-movement related activity with changes at the symbol switch. Using standard sterile surgical techniques and endotracheal isoflurane general anesthesia, we implanted 10–15 titanium screws in the monkeys’ skull and used them to anchor an acrylic cap in which we placed a head-fixing device and the recording chamber. We left the bone intact and made 3 mm burr holes through which we then could insert the electrodes. We used two recording cylinders, on the left hemisphere of each monkey.

### Task

#### Two-alternative forced-choice discrimination task

The task began with the monkeys grasping two bars, one with each hand, after which a white fixation point appeared for 800 ms. The monkeys were not constrained to fixate at the fixation point, but they inevitably did so. Then one of a pair of symbols appeared, briefly for 100 ms in some sessions or until the monkey initiated a hand response in other sessions, at the center of gaze. One symbol signaled the monkey to release the left bar and the other to release the right bar. We rewarded the monkeys with a drop of juice for releasing the hand associated with that symbol. We did not punish the monkeys for errors. We overtrained the monkeys on one set of symbol-hand association (Green Square for left hand and Pink Square for right hand). In the visuomotor associative learning version of the task (**Fig 1**), we began every recording session by presenting the monkeys with the same over-trained symbol pair (overtrained condition), and after a number of trials, switched the symbol pair to two fractal symbols, which the monkey had never seen before (novel condition), and did not have colors matching the overtrained symbols. Over 20 to 40 trials, the monkey gradually learned which symbol was associated with which hand. The manipulanda remained the same throughout the task. In the manipulanda change task (**Fig 2**), we began every recording session by presenting the monkeys with the same over-trained symbol pair and bar manipulanda, and after a number of trials, switched the bar manipulanda to dowel manipulanda. The visuomotor association remained the same throughout the task.

### Data collection

#### Single unit recording recording

We introduced glass-coated tungsten electrodes with an impedance of 0.8-1.2 MOhms (FHC) into the left mid-lateral cerebellum of monkeys every day that we recorded. We passed the raw electrode signal was through a FHC Neurocraft head stage, and amplifier, and filtered through a Krohn-Hite filter (bandpass: lowpass 300 Hz to highpass 10 kHz Butterworth), then through a Micro 1401 system, CED electronics. We used REX-VEX system coupled with spike2, CED electronics for event and neural data acquisition. We verified all recordings off-line to ensure that we had isolated Purkinje cells (**Fig 1B, S2**) and that the spike waveforms had not changed throughout the course of each experiment (**Fig S4**). We identified cerebellar Purkinje cells online by the presence of complex spikes **(Fig. 1B)**, and offline by the i) spike waveforms, ii) a pause in simple spike after a complex spike **(Fig S2A)**, and iii) the simple spike interspike interval distribution (*17*) (**Fig S2B**).

#### Hand tracking

We either painted a spot on the monkeys’ right hand with a UV-blacklight reactive paint (Neon Glow Blacklight Body Paint) prior to every session or tattooed the right hand with a spot of UV Black light tattoo ink (Millennium Mom’s Nuclear UV Blacklight Tattoo Ink). We used a 5W DC converted UV black light illuminator to shine light on the spot. Then we used a high speed (250 fps) camera (Edmund Optics), mechanically fixed to the primate chair, to capture a video sequence of the hand movement while the monkeys performed the tasks.

### Data Analysis

#### Preference indexes

We defined hand preference index (HPI) and symbol preference index (SPI) as the normalized difference in spiking activity between left and right hands or the two symbols presented during the over-trained task.

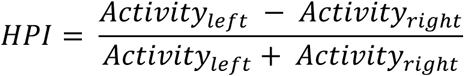

similarly,

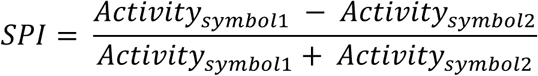

These indexes range from −1 to 1.

#### Hand movement data analysis

We used the track mate feature (*18, 19*) and custom written software in MATLAB to semi-manually track the fluorescent paint spot painted on the monkey’s hand. First, we used a downsampled LoG (Laplacian of Gaussian) filter, usually with an estimated blob diameter of 20 and a threshold of 20, to detect the fluorescent dot on the hand. Then we chose a range of threshold manually to detect the dot with considerable fidelity. We then tracked the spot using a nearest neighbor search approach. Briefly, the search algorithm relies on the KD-tree technique whereby the spots in the target frame are searched for the nearest neighbor of each spot in the source frame. If the spots found are closer than the maximal allowed distance (15 pixels), a link between the two spots is created and the process is repeated. We further confirmed the tracked path by using a linear assignment problem (LAP) tracker (*20*), allowing for gap filling. We manually removed spuriously detected spots from each frame in the tracks, during post processing. We then analyzed the tracks in MATLAB using custom written software. We smoothed the raw trajectories by using low pass moving filter with filter coefficients equal to the reciprocal of the span. We aligned all trajectories to the first instance of hand release from the bars. We excluded hand trajectory outliers from our data set if we could not reliably trace the trajectories.

### Method to detect changes in neural activity pattern

We developed a novel method to quantify the change in activity pattern between two conditions A and B. Suppose condition A precedes condition B,. We compared the activity within the first condition (A) with activity across both conditions (A and B). Both A and B are 20 × n matrices with n being the length of the signal. First, we randomly sampled 10 trials each from the last 20 trials in condition A and the first 20 trials in the condition B and calculated the root mean squared (rms) distance between the mean activities:

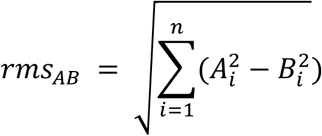

Where 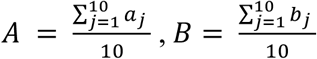 and n is the length of the signal

Also, a_j_ and b_j_ were sets of 10 trials chosen randomly from 20 trials.

We repeated this process 250 times to obtain a distribution of rms distances that compared the extent of change in across-condition activity profile between conditions A and B. To compare this distribution with the within-condition activity profile, we randomly sampled 10 trials twice without replacement, from condition A and repeated the same analysis to obtain another distribution of rms distances.

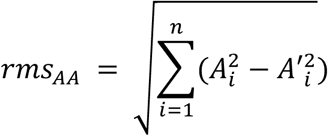

Where 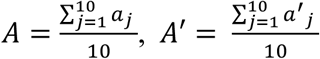 and n is the length of the signal

Also, a_j_ and a’_j_ were set of 10 trials chosen randomly from N and N-10 trials respectively, where N is the number of trials in condition A.

Next, we repeated this process 250 times to get a distribution of *rms_AA_* and *rms_AB_*. Finally, we compared means of both distributions for statistical difference. See **Fig S5** for validity of this method applied to Gaussian distributions and Poisson spike trains.

#### Statistics

To check if two independent distributions were significantly different from each other, we first performed a two-sided goodness of fit Lilliefors test, to test for the normality, then used an appropriate t-test; or else a non-parametric Wilcoxon ranksum test. All error bars and shading in this study, unless stated otherwise, are mean ± s.e.m.

